# *In vitro* culture of freshly isolated *Trypanosoma brucei brucei* bloodstream forms results in gene copy-number changes

**DOI:** 10.1101/2021.06.10.446608

**Authors:** Julius Mulindwa, Geofrey Ssentamu, Enock Matovu, Kevin Kamanyi Marucha, Francisco Aresta-Branco, Claudia Helbig, Christine Clayton

## Abstract

Most researchers who study unicellular eukaryotes work with an extremely limited number of laboratory-adapted isolates that were obtained from the field decades ago, but the effects of passage in laboratory rodents, and adaptation to *in vitro* culture, have been little studied. For example, the vast majority of studies of *Trypanosoma brucei* biology have concentrated on just two strains, Lister 427 and EATRO1125, which were taken from the field over half a century ago and have since have undergone innumerable passages in rodents and culture. We here describe two new *Trypanosoma brucei brucei* strains. MAK65 and MAK98, which have undergone only 3 rodent passages since isolation from Ugandan cattle. High-coverage sequencing revealed that adaptation of the parasites to culture was accompanied by changes in gene copy numbers. *T. brucei* has so far been considered to be uniformly diploid, but we also found trisomy of chromosome 5 not only in one Lister 427 culture, but also in the MAK98 field isolate. Trisomy of chromosome 6, and increased copies of other chromosome segments, were also seen in established cultured lines. The two new *T. brucei* strains should be useful to researchers interested in trypanosome differentiation and pathogenicity. Initial results suggested that the two strains have differing infection patterns in rodents. MAK65 is uniformly diploid and grew more reproducibly in bloodstream-form culture than MAK98.

## Introduction

*Trypanosoma brucei brucei* and related parasites infect humans and various domestic animals, and can be transmitted mechanically and venereally as well as by their definitive host, the Tsetse fly. *T. brucei gambiense* is the subspecies that causes West African sleeping sickness, while *T. brucei rhodesiense (*abbreviated to *T. rhodesiense)* causes East African sleeping sickness. *T. brucei brucei* (abbreviated to *T. brucei)* is found in cattle, but not humans; it differs from *T. rhodesiense* only in the absence of the *SRA* gene, which enables *T. b. rhodesiense* to survive in human plasma [1, 2]. Within Tsetse flies, *T. brucei* multiply as procyclic forms in the midgut, before migrating to the salivary glands, where sexual reproduction can occur, with meiosis and gamete formation, followed by gamete fusion [3, 4].

The *Tr. brucei* genome consists of eleven megabase-length chromosomes, which are generally diploid, and a variable number of “minichromosomes”. The parasite escapes the immune response by antigenic variation, expressing a single Variant Surface Glycoprotein (VSG). The expressed *VSG* gene is located at a telomere, and can be changed either through transcriptional switching or, more commonly, by genetic rearrangement. Every parasite has at least 2000 alternative *VSG* genes or pseudogenes, which are located in sub-telomeric arrays and on the minichromosomes [5, 6]. DNA contents vary up to 30% between *T. brucei* isolates [7]. Although some of this can be attributed to differing minichromosome contents [7], the lengths of the megabase chromosomes also differ substantially both within, and between, strains [7-9]. Variations in the numbers of *VSG* genes are to be expected, but there are also other differences. Some genes are arranged in multi-copy arrays, which facilitates high expression but also leaves the genes prone to homologous recombination. For example, the beta- and alpha-tubulin genes are present in an alternating array, and one study found fewer copies in *T. brucei gambiense* than in *T. brucei rhodesiense* or *T. brucei brucei* [7].

There is so far no evidence for mating in *T. b. gambiense*, but in *T. brucei rhodesiense* or *T. brucei brucei* the chromosome copy number might be expected to be made more uniform by meiosis and mating - although the sexual stage is not obligatory. Indeed, analysis of sequence data from 26 field isolates has so far revealed no evidence for aneuploidy [10]. Aneuploidy appears to possible, however: after mating experiments, although classical Mendelian inheritance is seen [11], triploid progeny also appear to be relatively common [12, 13]. The apparent uniform diploidy of salivarian trypanosomes contrasts with considerable aneuploidy in the stercorarian trypanosome *Trypanosoma cruzi* [14] and the more distantly related leishmanias (e.g. [15]).

In the past few decades, nearly all studies of *T. brucei* molecular and cellular biology have used just two strains: Lister 427, which was probably originally isolated in Uganda from a cow in 1956 (see http://tryps.rockefeller.edu/DocumentsGlobal/lineage_Lister427.pdf), and EATRO1125, which was isolated in Tanzania from a bushbuck in 1966 (see http://tryps.rockefeller.edu/DocumentsGlobal/lineage_antat1.pdf). These strains have been passaged innumerable times in rodents or culture. This selects for an accelerated growth rate, which presumably reflect changes in metabolism and cell-cycle regulation. Stumpy forms are growth-arrested bloodstream forms which are pre-adapted for differentiating into procyclic forms. Prolonged passage clearly selects for a diminished ability to enter cell-cycle arrest, and therefore loss of the “stumpy form” life-cycle stage. Since some genes that are required for survival as procyclic forms are not needed in the bloodstream, prolonged culture or rodent passage as bloodstream forms can also result in loss of the ability to differentiate into proliferation-competent procyclic forms. This has, for example, occurred for the Lister 427 bloodstream forms currently used for genetic manipulation. Although several *T. brucei* strains, including EATRO1125, have been maintained in such a way as to preserve their differentiation capacity, we do not know whether their differentiation pathways are identical to those found in natural populations.

Another problem is that cultured Lister 427 trypanosomes have been maintained for decades in separate labs. As a consequence, the parasites that we study now are likely to show considerable differences in sequence and regulation compared with their ancestors, and also between labs. This will have been exacerbated by multiple cloning steps that have occurred during the selection of lines suitable for genetic manipulation. Analysis of gene copy numbers indeed suggested that in comparison with recently-isolated *T. rhodesiense*, common “lab” strains of *T. brucei* had varying expansions in multicopy gene arrays encoding proteins required for rapid cell division, and also differed from each other [16].

In this paper we set out to establish new *T. b. brucei* strains that have undergone minimal passage since field isolation in order to expand the repertoire of trypanosomes available for lab investigation. We describe the infection and culture characteristics of two new Ugandan *T. b. brucei* strains that have very different disease profiles in mice. Available stocks of these parasites have undergone only 3 mouse passages since their isolation from cattle. These lines should be useful for analyses of factors governing *T. brucei* disease course, tissue distribution, and differentiation. In addition, we aimed to find out what happens when *T. brucei* are adapted to bloodstream-form culture. The growth of the new trypanosomes was slower than of standard lab lines but genome changes were rapid, with gene copy number changes after only a few weeks.

## Results and discussion

### *In vivo* growth of two new *T. b. brucei* isolates

*T. b. brucei* MAK65 was isolated from a cow in Banya parish, Apac district on February 1st, 2016, while *T. b. brucei* MAK98 was isolated from the same place on July 30th, 2016 (Figure 1). The identity as *T. brucei brucei* was confirmed by the presence of a 480bp band PCR of the rRNA internal transcribed spacer [17], and absence of the SRA gene [18] (S1 text). The strains were shown to be different by microsatellite typing [19] (S1 text). The frozen cow blood was passaged once through mice in order to make stabilates for further use.

**Figure 1.**
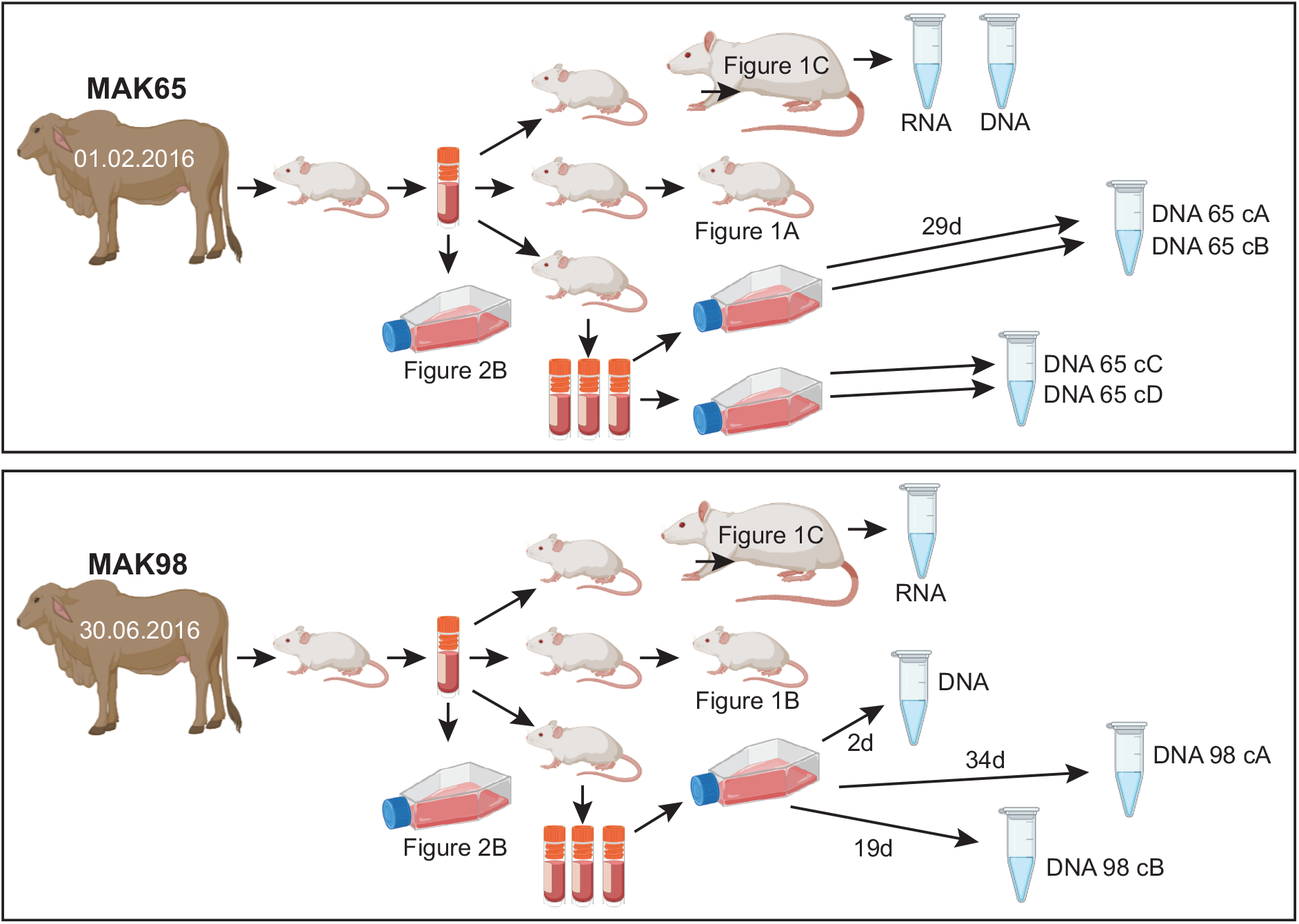
Parasite histories and culture. The images are from biorender (https://app.biorender.com)

To characterize the two isolates, we first infected 8-week-old swiss mice with 1000 parasites each (Figure 1). Parasitaemias are shown in Figure 2A and 2B. Results suggested that MAK65 might be less virulent than MAK98. We also infected rats with 5000 parasites, after a single previous mouse passage to avoid artefacts due to differences in stabilate viability (Figure 1). Parasitaemias were again lower for MAK65. We harvested the parasites 5 or 6 days after infection (Fig 2C). We had hoped that the populations harvested earlier in infection would show low expression of stumpy-form markers, and would therefore be suitable for characterization of long-slender-form transcriptomes. On the contrary, the proportion of stumpy forms (as judged by PAD1 staining [20]) was highest for MAK65 parasites at low parasitaemia (Fig 2C, Supplementary Figure 1A). Conversely, in the rat with 50-times higher MAK98 parasitaemia, no stumpy forms were detected (MAK98B) even though cell proliferation had clearly slowed. It has been proposed that development of stumpy forms is primarily a mechanism to limit virulence [21]. From these results it seems possible that MAK65 has lower virulence in mice than MAK98 because MAK65 undergoes stumpy-form differentiation at a lower density. However, we cannot rule out the possibility that MAK65 has higher densities in tissues such as fat or skin. Also, it is conceivable that the inocula differed in infectivity; this would have happened if one had a higher proportion of long slender parasites. To confirm the differences in virulence and - if present - determine its basis it will be necessary to examine several more infections, and to characterize parasites in tissues as well as the blood.

**Figure 2.**
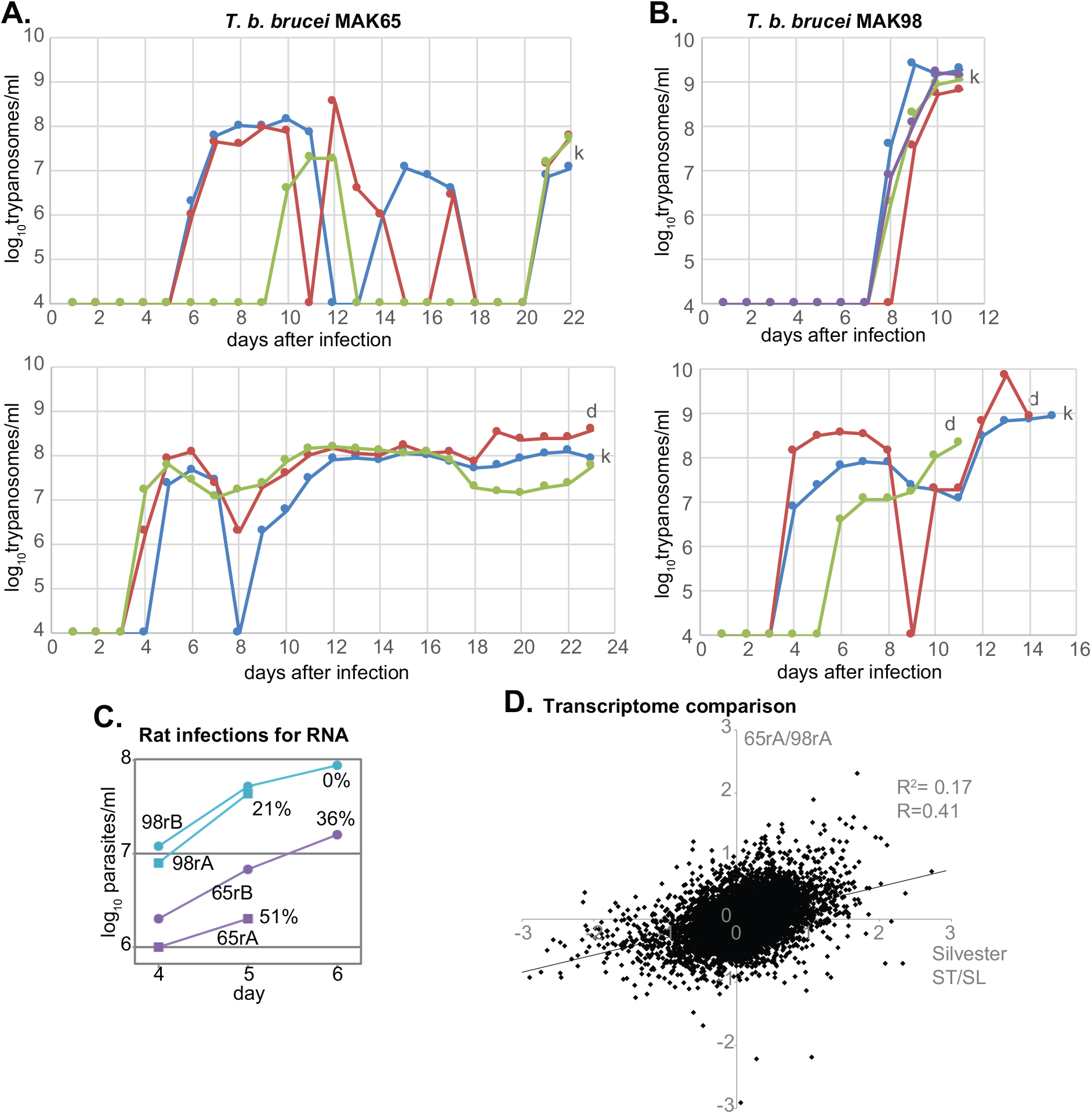
Growth of MAK65 and MAK98 strains in rodents. A. Parasitaemias after infection of three or four mice with 1,000 strain MAK65 parasites. “d” indicates death and “k” indicates killing because of clear symptoms. The y axis scale is the log10 of the parasitaeimia. B. Parasitaemias after infection of 3 mice with 1,000 strain MAK98 parasites. C. Growth of both strains in rats before harvest for RNA purification (rA= rat A, rB = rat B). The percentage of PAD1 positive cells at the time of harvest is indicated. D. The transcriptomes of population 65 (rat A) and 98 (rat A) parasites (panel C) were compared and the log2 ratio (65A/98A, Supplementary Table S1, sheet 1) is on the y-axis. The x-axis shows published stumpy/long slender results for trypanosomes grown in mice [22]. The correlation coefficients are from linear regression analysis using the log-transformed rations, and were generated in Microsoft Excel.

Transcriptome analysis (E-MTAB-9320, S1 Table) revealed that parasites of the MAK65-rat-A (65rA) population shown in Figure 2C had highest expression of mRNA encoding the procyclic-form surface protein GPEET procyclin as well as other procyclic-specific mRNAs such as those encoding trans-sialidase and enzymes of the citric acid cycle. Interestingly, although the parasites for MAK98-rat-B did not show PAD1 staining, the population with the lowest expression of stumpy-form markers was 98-rat-A. Comparison of the 65-rat-A and 98-rat-A transcriptomes with those of pure EATRO1125 long slender and stumpy forms [22] confirmed that MAK65-rat-A indeed had a more stumpy-form like expression pattern than 98A (Figure 2D). Although these were by no means pure populations, the correlation (R=0.44) was better than that obtained for *in vitro* stumpy-form differentiation of EATRO1125 [23] (S1 Fig. B, R=0.33). These results suggest that differentiation to stumpy forms in immune-competent rats resembles differentiation in immunosuppressed animals, although more biological replicates would be required to confirm this conclusion.

We did not compare our new transcriptomes directly with the previous data from EATRO1125 because differences in cell harvesting, RNA preparation and sequencing methodology would be likely to affect the outcome. Such differences are at least partially eliminated when ratios are compared. For example, we had already obtained some evidence that poly(A) selection selects against long mRNAs, but this was based on comparison of different RNA samples [24]. We now confirmed the length effect. Starting with the same initial total RNA samples, we made mRNA by oligo d(T) selection, or by depleting rRNA using RNase H and complementary oligonucleotides. We had previously demonstrated, by comparing with total RNA, that the latter procedure has no effect on the transcriptome [25]. Analysis of the transcriptomes clearly showed that oligo d(T) selection causes loss of longer RNAs (S1 Fig C).

We also sequenced genomic DNA from the two new strains. The initial populations that were sequenced were MAK65 from rats, and MAK98 cultured for 2 days (see below) (Figure 1). We obtained Oxford nanopore reads (E-MTAB-9318) as well as 72-nt paired-end Illumina reads at over 100-fold coverage (E-MTAB-9759). In combination these should allow genome assembly. To minimize manipulation, we did not clone these cells at any stage. Although *T. brucei* populations are to some extent clonal [26, 27], we cannot rule out the possibility of mixed infections, and within-population variation is also expected.

### *In vitro* growth of the two new *T. b. brucei* isolates

To assess the abilities of the two new strains to grow *in vitro*, we placed trypanosomes directly from mouse blood stabilates into HMI-9 medium and followed cell numbers, diluting them regularly to prevent densities in excess of 1.5×10^6^/ml. Both isolates grew rapidly from stabilate but after 1-2 days, the growth slowed markedly and became intermittent (S2 Fig A, B) with variable doubling times of up to 2 days. As expected, there were clear deleterious effects when the density accidentally exceeded 2×10^6^/ml. To find out whether the growth in culture was reproducible, new stabilates were thawed and placed into culture. Full cumulative curves for all samples that were subsequently used for genome characterization are in Fig 3, and details illustrating the dilutions are in S2 Fig C-I. The MAK65 cells again adapted to culture quite readily, with an initial division time of about 17h which shortened to about 11 h after 5 days (S2 Fig C). (This might indicate a selection bottleneck, with some cells dying or growing slowly, while others grow more readily.) During culture for a further 2 weeks, growth remained somewhat erratic, with clear deleterious effects when the density accidentally exceeded 1.0×10^6^/ml, but the division times progressively decreased to about 7h (S2 Fig. F; cultures MAK65cA, MAK65cB). Similar observations were made for two additional cultures initiated from a new stabilate (S2 Fig. G; cultures MAK65cC, MAK65cD).

**Figure 3.**
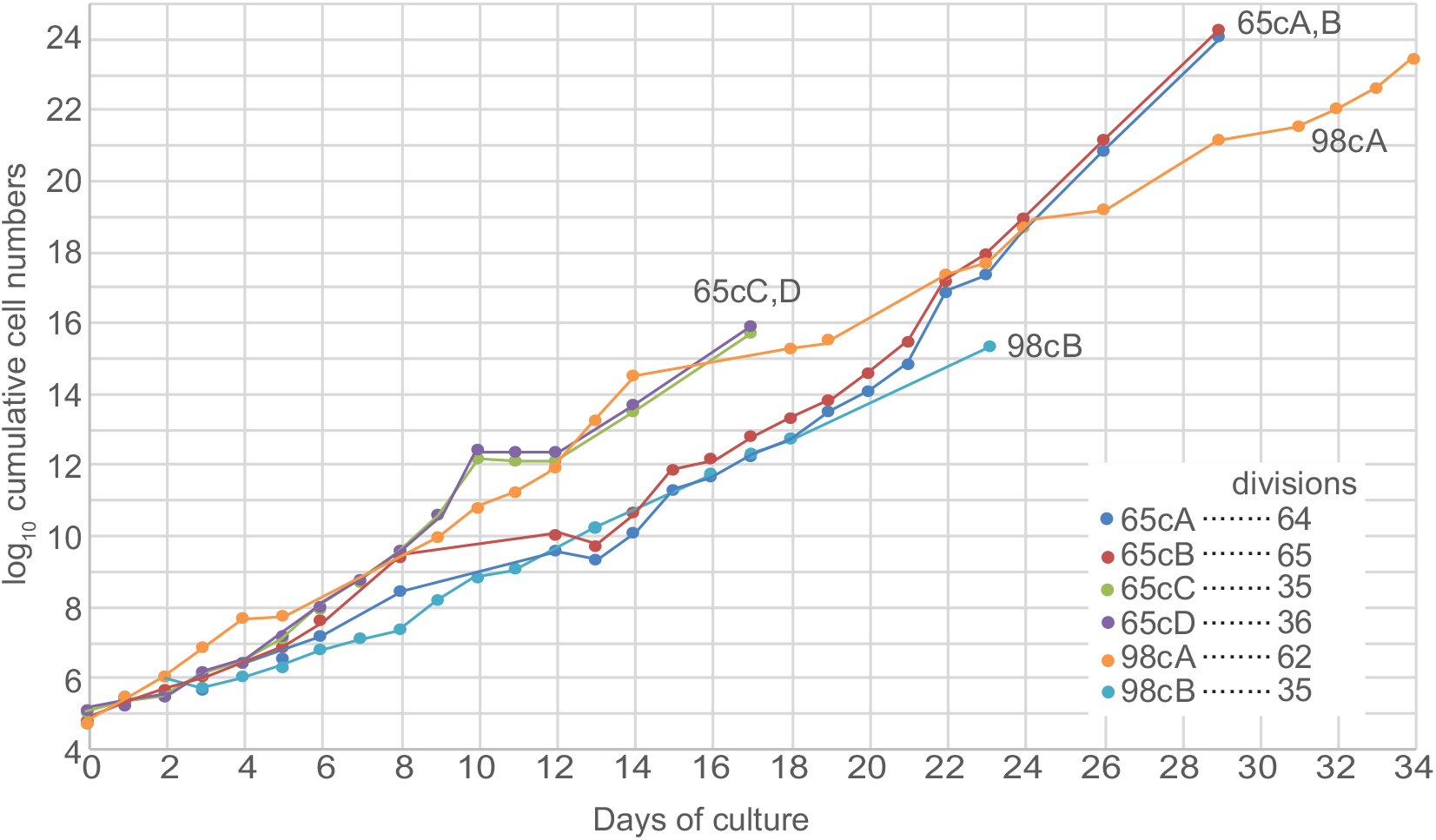
Growth in culture. Cumulative cell counts for all cultures that were used for DNA preparation. Full details of these cultures with linear-scale plots are in S2 Fig panels C-I.

For MAK98, over the first 24h the division time was 8h (S2 Fig D; this is the culture that was used for genomic sequencing). Cells were then divided into 10 replicate cultures with different starting densities varying from 6×10^3^ /ml to 2×10^5^/ml. Over the next 24h the division time was 17±3h irrespective of the starting density (S2 Fig G) and the maximum density that was subsequently obtained was 1.5 ×10^6^/ml. Several additional attempts to culture MAK98 from frozen blood stabilates failed. Although this might be an artefact - a property of the particular stabilates used - the subsequent slow growth of MAK98 in culture suggests that might be intrinsically less culture-adaptable. For prolonged growth, we first continued the MAK98 culture shown in S2 Fig D, giving MAK98 culture A; this grew erratically with an overall division time of 13h despite never exceeding densities of 5×10^5^/ml (S2 Fig. H). Later, we initiated a new culture from a frozen stock made from the initial two-day culture; this resulted in culture MAK98 culture B, which after a further week achieved a division time of 11h (S2 Fig. I). A comparison of all cultures together (Fig. 3) illustrates the slower proliferation of both MAK98 cultures relative to MAK65.

Very surprisingly, neither strain survived in medium containing 1.1% methyl cellulose [28] which we were concurrently using for passage of EATRO1125 bloodstream forms. Results of preliminary experiments suggested that it was possible to obtain procyclic forms by cultivating the cells with cis aconitate at 37°C for 24h, then placing them in procyclic-form medium (SDM79) at 27°C, but this has not been investigated in detail.

For future genetic manipulation attempts, the MAK65 line appears preferable because it is uniformly diploid (see below), and appears to be easier to culture than MAK98.

### *In vitro* growth affects gene copy numbers

To find out how culture affects the genome, we sequenced MAK65 that had been cultured for 4 weeks (65 culture A, 65 culture B, E-MTAB-10457) and 2 weeks (65 culture C, 65 culture D, E-MTAB-10457); and MAK98 that had been cultured for 7 weeks (MAK98 culture A, E-MTAB-10466) and 3 weeks (MAK98 culture B, E-MTAB-10457).

Given the uncloned nature of the starting populations we expected mainly to see selection of parasites with better abilities to grow in culture, and perhaps of new mutations. Once the starting genomes have been assembled, it will be possible to do detailed analyses of single nucleotide polymorphisms that were selected. We had previously found differences in numbers of multi-copy genes between lab-adapted trypanosomes and *T. b. rhodesiense* isolated from patients [16, 24], but these analyses were constrained by the limited number of trypanosome populations available and in particular, the absence of data for un-passaged versions of the lab-adapted strains. We therefore measured variations in gene copy number in the new data. To do this we used a published list of open reading frames that contains all single-copy genes, plus just one representative of genes that are repeated (>90% identity) in the *T. brucei* TREU927 genome, with addition of a few ribosomal protein genes that had been omitted on the basis of short length [29]. The read densities over each open reading frame (reads per million per kilobase, RPKM) were calculated (S2 Table). We initially then divided the RPKMs obtained by the modal RPKM value, on the assumption that most genes are present in a single copy in the haploid genome. The distributions obtained were then examined, and the denominator was adjusted to give a peak value of approximately 2 copies per diploid genome.

First, we re-analysed the genomes of trypanosomes we maintain routinely in the laboratory, comparing them with the previous previously-characterized *T. b. rhodesiense* strains, and the reference “genome” strain, TREU927. We looked at data from three long-term-cultured bloodstream-form lines (two Lister 427, one EATRO1125) and one Lister 427 procyclic-form culture; all had been cloned at least ten years previously, after genetic manipulation, and had been cultured intermittently since. Figure 4 shows the copy number distribution for the majority of genes - those with 0.5 - 2 copies per haploid genome; and copy-numbers across the genome are shown in Figure 5. The “genome” reference strain TREU927 was initially chosen because the genome size is relatively small relative to other strains, and because it is differentiation-competent. The source of the DNA for TREU927 sequencing is not clear from the publication: it may have been derived from procyclic forms that had recently-differentiated from blood parasites [8], but in that period isolation from rodent blood was more common. All TREU927 chromosomes were diploid, though the copy-number distribution for individual genes was broader than for the un-passaged *T. rhodesiense* (Figure 4A, Figure 5A). We discovered that our EATRO1125 culture has three copies of small regions of chromosomes 1 and 2, and has haploid loss of a small segment at the end of chromosome 11 (Figure 5B).

**Figure 4.**
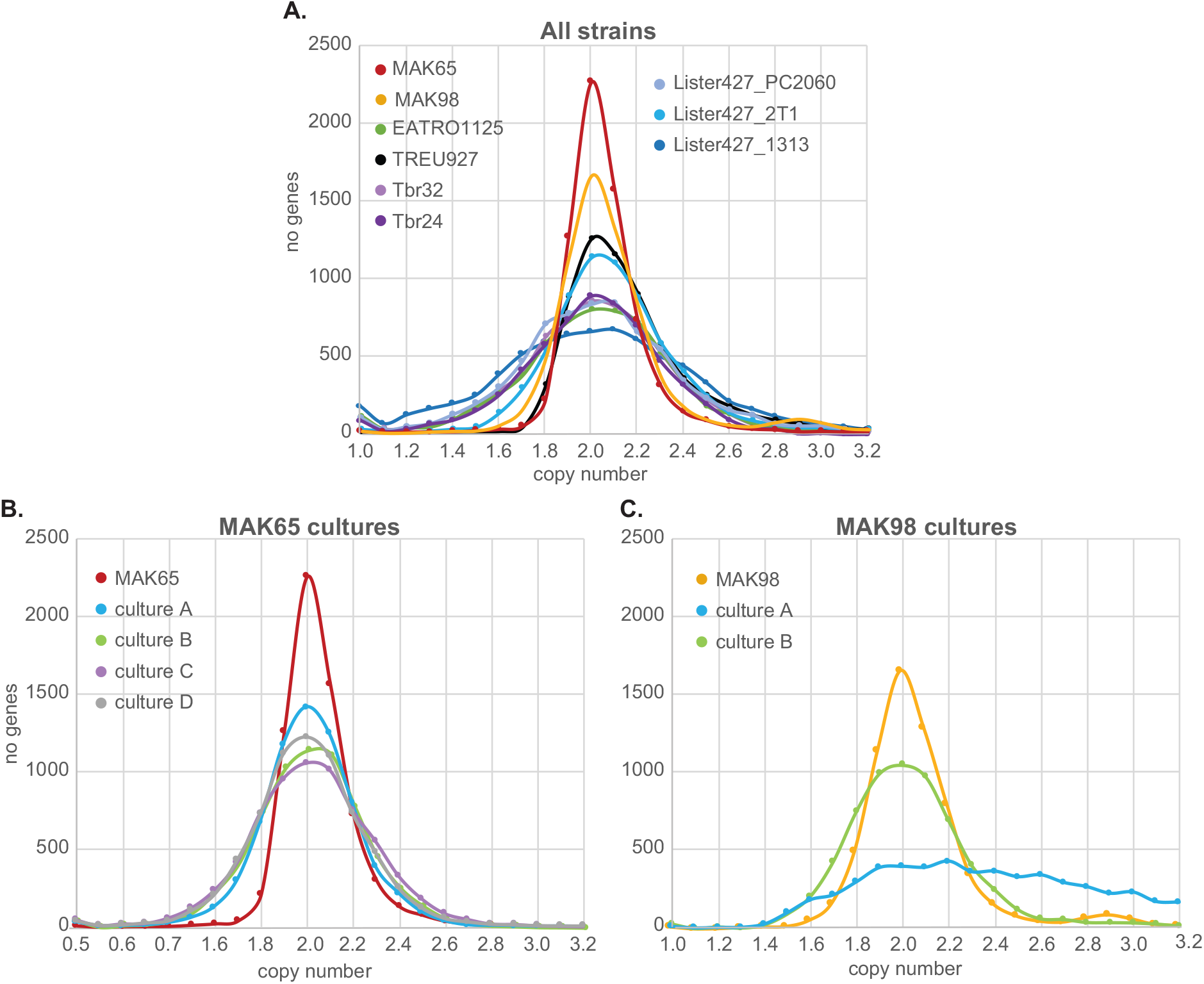
Copy number distributions for different trypanosome strains and cultures. After genome sequencing all reads were allowed to align 20 times. For a set of unique genes, containing one representative each for each set of repeated genes, reads per million reads were calculated. Copy numbers for each gene were then calculated based on the assumption that most genes are present once in every haploid genome (or twice per diploid genome). The numbers of genes with copy numbers between 1 and 4 are plotted here for the strains and cultures indicated. A. *T. rhodesiense*, TREU927, EATRO1125 and Lister 427. B. MAK65 from rats and culture C. MAK98 from rats and culture.

**Figure 5.**
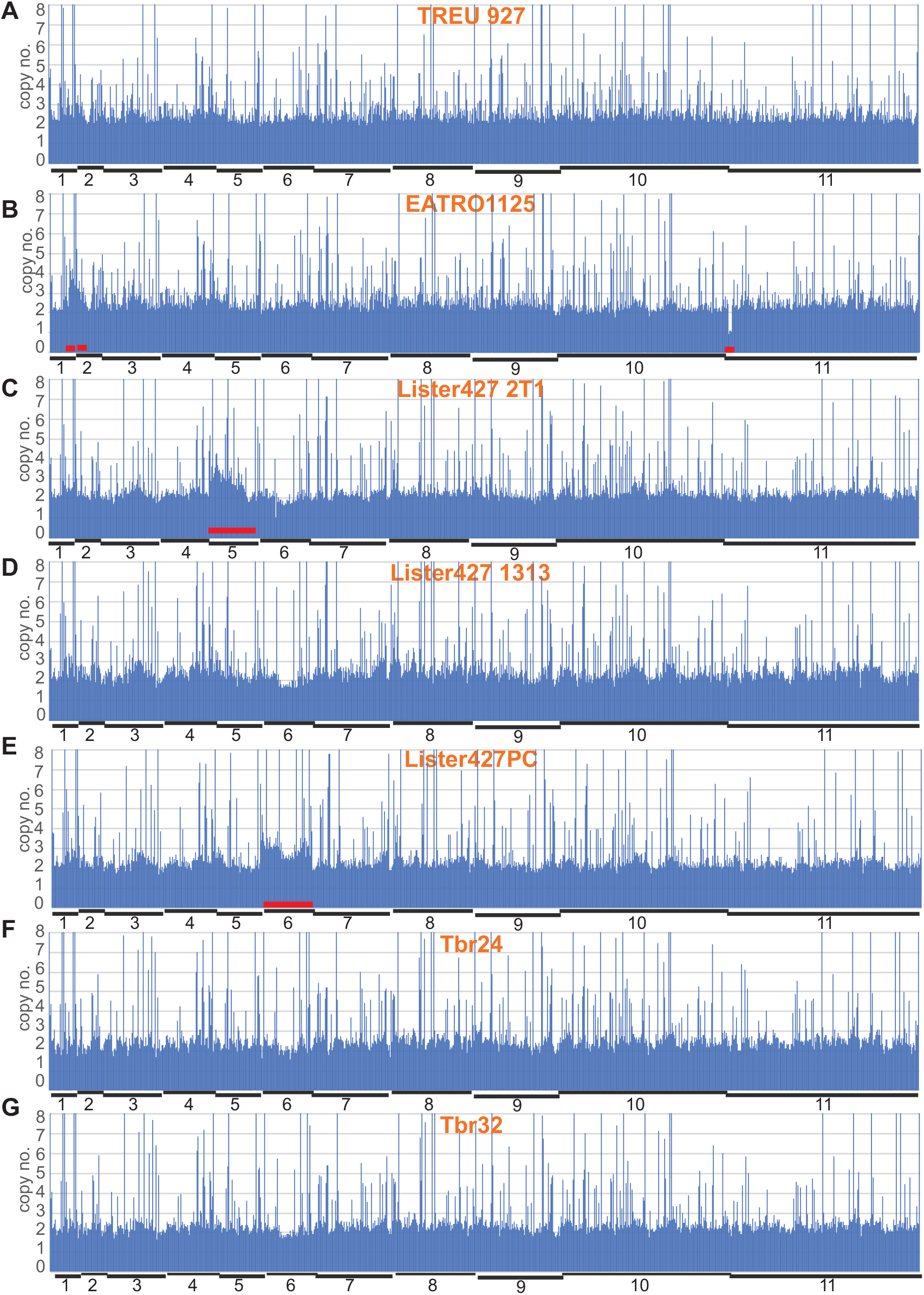
Gene copy numbers of standard strains and a *T. rhodesiense* isolate, plotted across the genome. Copy numbers for each unique open reading frame were determined as in Figure 3. The unique genes were then placed in order along the chromosomes (S3 Table sheet 3) and their copy numbers plotted with one bar for each unique gene. The approximate chromosome boundaries are indicated on the plot; columns exceeding 8 copies are truncated at the top of each panel. Red bars indicate departures from diploidy. The nature of the sample is indicated on each panel. A: TREU927 (genome strain); B: Cultured EATRO1125 bloodstream forms expressing the tet repressor; C: Lister 427 2T1 bloodstream forms expressing the tet repressor and T7 polymerase (427_2T1) [45]; D: Lister 427 bloodstream forms expressing the tet repressor (427_1313) [31]; E: Lister 427 procyclic forms (427_PC2060) expressing the tet repressor; F, G: *T. b. rhodesiense* from humans, passaged 3 times in rodents [16].

These discrepancies most likely represent intra-chromosomal duplications and deletions. A duplication in a ∼200kb segment of chromosome 2 was also reported for some field isolates [10]. The “2T1” Lister 427 line, which contains a “landing pad” that allows targeting of plasmids to a specific rRNA spacer [30] showed numerous copy-number differences from Lister 427 carrying pHD1313 [31], including apparent trisomy of chromosome 5 (Figure 4C). The broadest copy-number distribution was for a bloodstream-form Lister 427 line containing pHD1313 (Figure 4A, 5D); while our procyclic-form tet-repressor-expressing Lister 427 line was trisomic for chromosome 6 (Figure 5E). A recent preprint describing copy numbers (analysed using a slightly different approach) reported similar changes for chromosome 5 (bloodstream form) and 6 (procyclic form) [5]. (The histories of the different cultured lines are complicated and these duplications may not be independent events.) In the preprint trisomies in chromosomes 2 and 7 were also mentioned. Trisomies would compromise attempts at homozygous gene deletion, unless Crispr-Cas is used. These results highlight the fact that strains with the same name, but grown for protracted periods in different laboratories, may have diverged considerably. *T. rhodesiense* derived from patients in a single sleeping-sickness focus, with minimal rodent passage [16], were uniformly diploid (Figure 4A and Figure 5F,G).

We next compared the genomes of the new strains with the previous ones. For both strains, most genes were diploid (Figure 4B, C, Figure 6 A, D) but MAK98 was again triploid for chromosome 5 (Figure 6 D). This is, to our knowledge, the first time that trisomy has been reported for a field isolate. Notably, the trisomy was also reflected in the transcriptome: comparison of MAK98 and MAK65 mRNA abundances revealed clear over-representation of mRNAs spanning chromosome 5 (Figure 6G, H). Thus changes in gene copy number are generally likely to be reflected at the mRNA level - although additional controls may affect protein stability or modification. Consentino et al. [5] also concluded that changes in ploidy were not compensated at the level of mRNA abundance..

**Figure 6.**
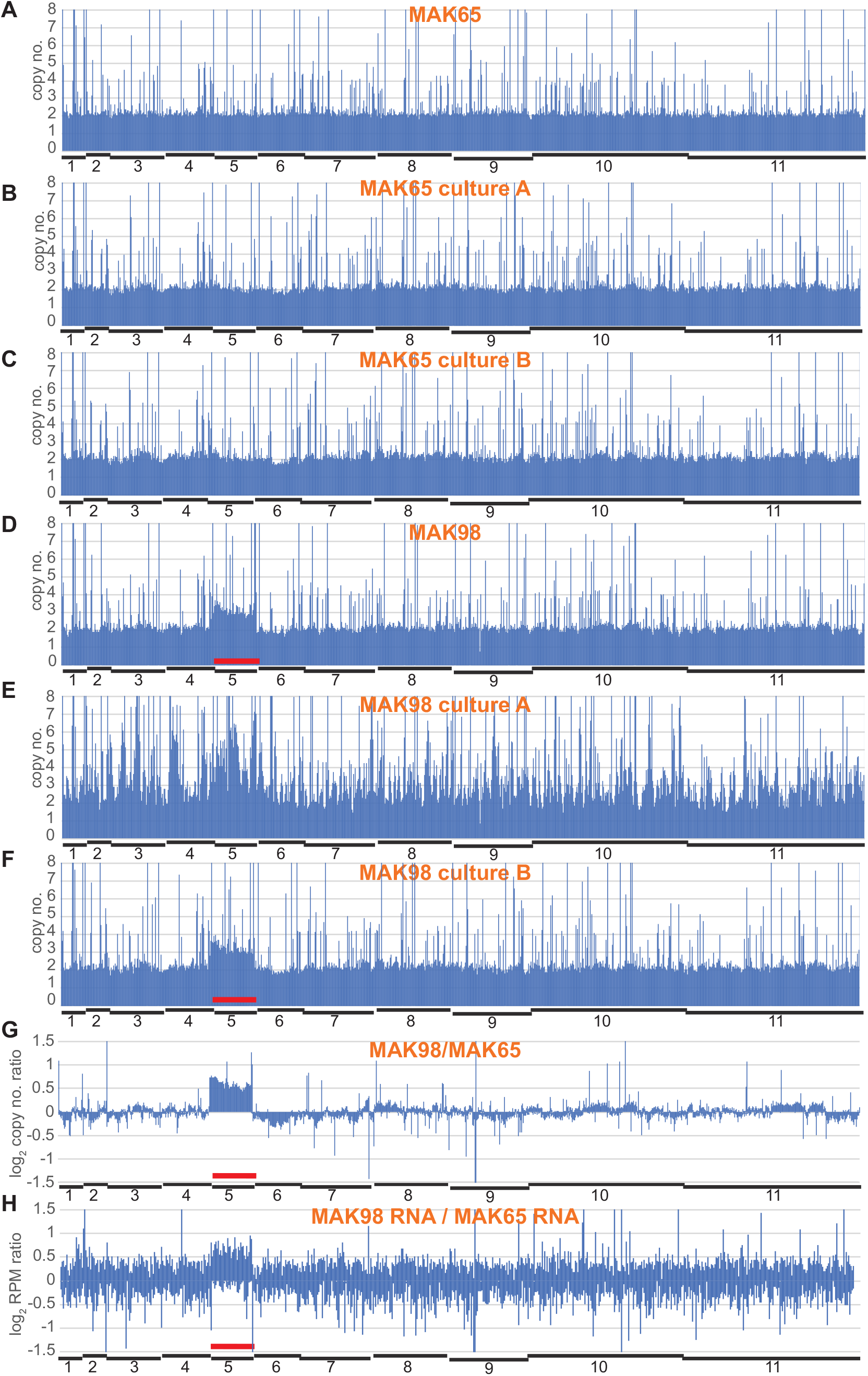
Gene copy numbers for original and cultured MAK65 and MAK98 A-F: Copy numbers plotted across the genome, details as in Figure 4. G: The average read densities for DNA from rat-grown MAK98 were divided by those for MAK65 and plotted across the genome (panel D / panel A). The results are on a log scale. H: The average read densities for RNA from rat-grown MAK98 were divided by those for MAK65 and plotted across the genome. The results are on a log scale. Many differences are likely to be due to the differentiation state (Figure 1 C, D) but MAK98 shows clear RNA over-representation from chromosome 5.

It was interesting that the *T. rhodesiense* strains from human patients showed somewhat more heterogeneity in gene copy-numbers than the two cattle-derived *T. brucei* (Figure 4A). The difference is unlikely to reflect geographical origin since the *T. rhodesiense* isolates were from Lwala, in Kaberamaido district, which is only about 100 km from Banya (Apac district) and the dates of isolation were less than 5 years apart. The difference is also unlikely to have been caused by growth in different hosts since cattle are a reservoir for *T. rhodesiense*, and *T. rhodesiense* almost certainly undergoes genetic exchange with *T. brucei* [32].

Surprisingly, just two weeks of culture was sufficient to broaden the copy-number distribution somewhat for both MAK65 and MAK98 (Figure 4 C, D). We do not know precisely how many times these cells had divided: the minimum number of generations is 35 (Figure 3), but the actual number is likely to be higher because some cells in the population may have failed to divide at various times during the culture period. Since the populations were not cloned, it is likely that culture resulted in selection of parasite subpopulations with either deletions, or duplications of different chromosome regions, as well as with particular single-nucleotide polymorphisms (SNPs). Duplications of individual single-copy genes might occur via homologous recombination involving low complexity sequences in untranslated regions. Despite some overall change, gene copy numbers of all MAK65 cultures looked broadly similar to the source population; examples are in Figure 6B and C. One of the MAK98 cultures - MAK98B - also looked similar to the starting population although some changes were evident (Figure 4C, 6F). In contrast, the MAK98A culture, which had been cultured for longer, had an exceptionally broad gene copy-number distribution profile, suggesting that an unusual number of genes in the population had undergone duplication or deletion (Figure 4C). Many small segments had changed in copy number throughout the genome (Figure 6E), with some chromosome segments present in triploid, tetraploid or more copies. We do not know whether the additional gene copies are internal chromosome duplications, or extra small chromosomes. This result, combined with the slow growth rate, hints that in addition to selection for particular variants, the parasites might have suffered defects in chromosome replication and/or segregation.

### *In vitro* growth may select for increased copies of specific genes

Finally, we looked to see whether there were any consistent changes in copy numbers after culture adaptation. First, we examined changes that happened in the MAK65 cultures and MAK98 culture B. (At this point we did not consider MAK98 culture A because it appeared to be so severely compromised.) 29 genes showed an increase of at least one copy for both MAK98 and MAK65 and 20 a similar decrease (Supplementary Table S2, sheet 1). Increases were seen for genes encoding the core histones, the major cytosolic chaperone HSP70, two translation elongation factors, three paraflagellar rod proteins, glycerol kinase, and the cyclin F box proteins CFB1 and CFB2. (The last two share sequence and scrutiny of the read coverage did not enable us to work out whether just one of the two genes was affected.) Examples are in Figure 7 A-H. Decreases were seen for rRNA genes (Figure 7I). and PIP39. The change in rRNA genes is surprising since rRNA is needed for rapid growth. PIP39 promotes differentiation of stumpy forms to procyclic forms [33] so the advantage of selection for decreased expression is not obvious. In order to find out whether these increases were also seen in Lister 427 or EATRO1125, we compared them to the average for the four *T. rhodesiense* genomes, which are likely to be closely related so cannot be considered independently. None of the decreases in gene copy number was reproducible. For example, the rRNA copy numbers in MAK65 and MAK98 had simply started out rather high in the initial populations, then decreased during culture to the numbers already present in Lister427 cultures (Figure 7I).

**Figure 7.**
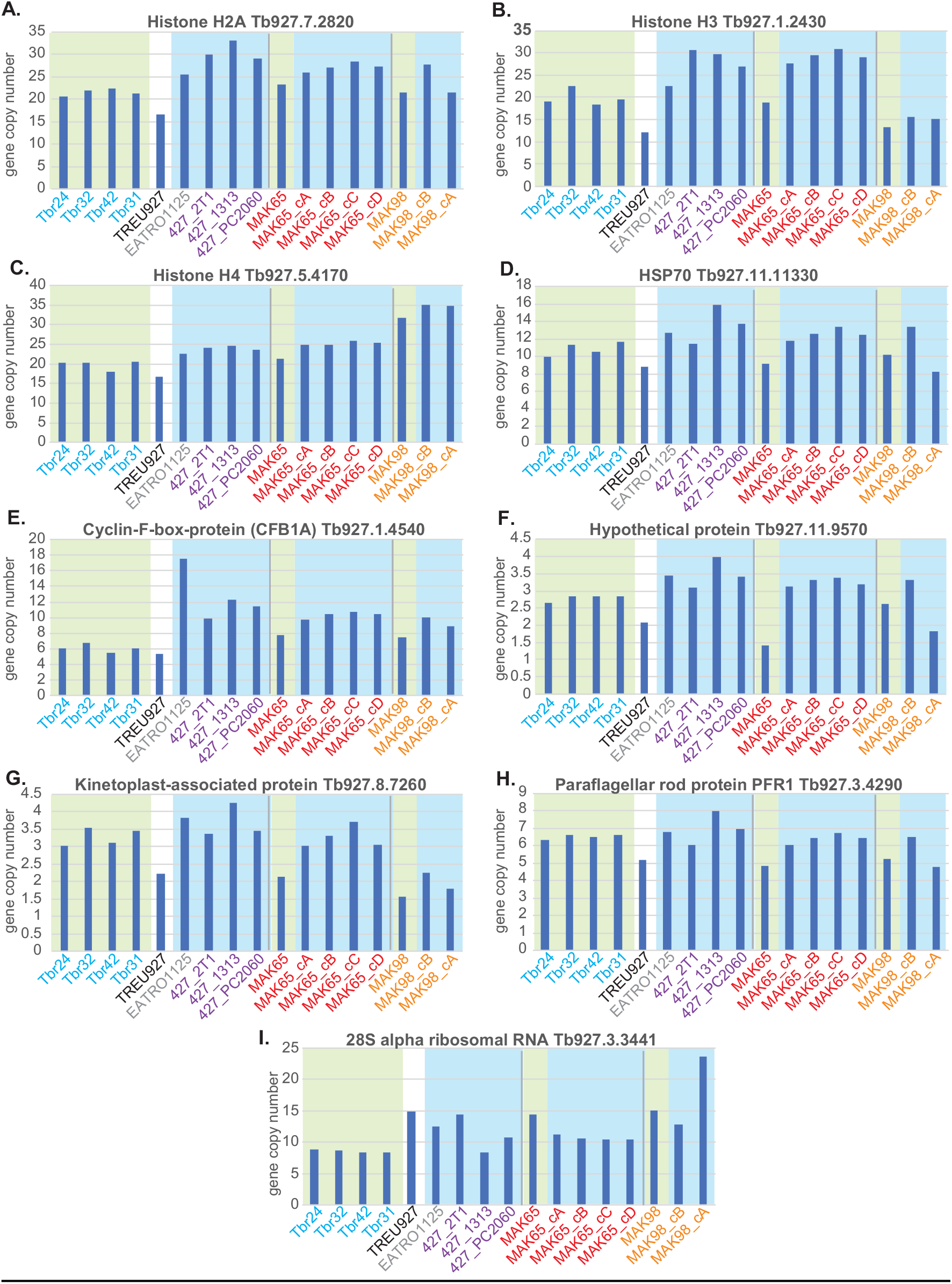
Copy numbers of selected genes that appeared to be influenced by *in vitro* culture. The gene products and numbers are shown above each graph; the y-axis shows the haploid gene copy number taken from S3 Table, sheet 1. Columns with green underlay are for non-culture-selected parasites, and those with blue underlay were culture-selected. Colours indicate the origin. The four *T. b. rhodesiense* isolates (Tbr) are from independent patients but have been assigned the same colour because they were all taken from a single clinic. The same colours for underlay and text are used in Supplementary table 2. Panels A-I are different genes as indicated.

With the small numbers of isolates and cultures examined here, and the extensive variation in initial copy numbers, none of the increases seen was statistically significant. Nevertheless, it was interesting that genes encoding the core histones H2A, H3 and H4 were in excess in cultures for all comparisons (Supplementary Table S2, sheet 1 and Figure 7, A-C). (H2A and H2B are sufficiently similar that only one of the two was considered.) These are all present as tandem repeats, which facilitates inter-and intra-chromosomal recombination; but there was clear specificity because many repeated genes showed no changes (S2 Table, sheet 3). Selection for histone gene amplification can be explained because faster-growing cells have a selective advantage. HSP70 genes also were higher in the cultures (Figure 7D); this might be needed to respond to culture-specific stresses. The CFB1/CFB2 cluster was also amplified in all cultures (Figure 7E). The function of CFB1 is unknown [34], while CFB2 binds to and stabilizes the mRNA encoding the variant surface glycoprotein [35]. Figure 7 F-H includes three other genes that showed increased numbers in most comparisons.

A detailed alignment of the region that includes *HSP70* is shown in S3 Fig. It compares one MAK65 culture (culture A) with the starting population. The *HSP70* gene (Tb927.11.11330) is arranged as tandem repeats, but is present only once in the TREU927 reference genome, because the (approximately 8) repeats were eliminated during genome assembly. The alignment for the starting population reveals differences between the homologous chromosomes (changes, relative to TREU927, seen in only half of the reads) and additional heterogeneity.

Comparing *HSP70* reads with those over the surrounding region, it is clear that in the culture, amplification occurred within the tandem repeat, without affecting neighboring genes. The alignment also shows that in this culture some SNPs and deletions are more abundant than in the starting population.

### Outlook

Our results show that cultured lines from the same original trypanosome isolate can have differences in karyotype, and that gene copy numbers and even chromosome ploidies in cell lines that are in routine use are very likely to differ between labs and clones, even if the parasites had the same origin. Perhaps we should not be surprised, therefore, if some results are not consistent between laboratories. It is perhaps also worth checking ploidies when thinking about gene knock-outs.

Initial growth of our two field-isolated parasite isolates in culture was relatively slow, but usually began to speed up after a few weeks. We do not know what is “missing” from the culture medium, relative to mammalian blood and tissue fluids. The observed slow proliferation could reflect slow overall metabolism in all of the parasites, but could also reflect a heterogeneous population, some growing quite fast, and others dying. The gene copy number changes seen after only a few weeks of culture adaptation might arise from selection of existing variants but we cannot rule out new recombination events. Our observations on copy number are the tip of the iceberg: a survey of just a single ∼10 kb region revealed possible selection for smaller insertions, deletions and point mutations (S4 Fig). To investigate this further it would be necessary to assemble the genomes - a non-trivial task [5] - and to study additional, newly-cloned, populations. As also noted by Cosentino et al., [5]. it would be really interesting to compare the genome of the original Lister 427 line [36] with those in routine use today. The results clearly raise the question of how much other cultured - or even rodent-adapted - organisms differ from their original counterparts, and to what extent the parasites that are studied in the laboratory are representative of those in the field.

*Leishmania* parasites have long been known to show a degree of aneuploidy, and to lose infectivity for mice if they are grown for too long as the promastigote form, which normally multiplies in the sandfly vector. A recent preprint described in great detail the effects of this culture on the *Leishmania donovani* genome [37]. The authors found copy number changes after only 20 generations, with progressively greater changes thereafter. In Leishmania, histone genes were not affected; instead, gene amplification predominantly affected genes involved in translation. They also observed deletions, sometimes of individual genes [37]. Although the affected genes may be different, the overall conclusion is the same: culture of trypanosomatid parasites rapidly selects for copy-number changes in specific genes.

Work on EATRO1125 and Lister427 has yielded innumerable insights into *T. brucei* biology, but we do not know to what extent this has been influenced by half a century in a laboratory setting. We here provide two new *T. brucei* strains for biological studies, with possible differences in rodent pathogenicity. Full genome assembly will enable a much more thorough comparison between the uncultured and cultured cell populations, as well as an assessment of the *VSG* gene repertoires of parasites subject to selection in the field. We did not clone the isolated parasites, or the cultures, so it would be interesting to compare several cloned lines both genetically and biologically. The courses of infection - including tissue distribution and capacity for differentiation to stumpy and procyclic forms, both *in vivo* and *in vitro*, could be investigated and compared with previous observations. Results so far suggest that MAK65 is an ideal model for such studies: it is diploid, readily generates stumpy forms, and can be cultured easily.

## Materials and Methods

### Trypanosome samples and infections

Peripheral blood (3-4ml) from cattle was collected into an EDTA tube (BD Vacutainer). An aliquot of the whole blood (600µl) was cryopreserved, a drop (10µl) was spotted on Whatman paper for PCR diagnosis. To determine the genus of the Trypanosome isolates, PCR was carried out on the Internal transcribed spacer (ITS-1) as described by Njiru et al 2004 [18].

To follow mouse infections, a stabilate was thawed and injected into a mouse. Once these parasitaemias had attained about 5×10^7^ trypanosomes/ml, 2-week-old inbred Swiss white mice were infected with 500 parasites each.

Parasites were counted by diluting 10µL of tail blood into 1mL of phosphate-saline-glucose, then counting in a haemocytometer. Thin blood smears were also prepared for immunofluorescence staining by fixed in methanol and permeabilizing with 0.2%TritonX-100. Mice were euthanized if obvious terminal symptoms were observed.

For RNA preparation, rats were infected with 5,000 parasites each. Parasites were followed using wet blood films. After the times shown in Figure 2C, blood (3-4mL) was collected by cardiac puncture and approximately 2.5ml drawn into a Paxgene tube. The Paxgene blood was incubated at room temperature for one hour and thereafter centrifuged at 5,000g for 10min. The supernatant was discarded, the pellet washed once with nuclease free water by centrifuging at 5,000g for 10min. The pellet was then resuspended in 1ml of Trifast reagent (Peqlab, GmbH) and transferred to 1.5ml microfuge tube for RNA preparation according to the manufacturer’s instructions. Parasite numbers in the 2.5 ml samples were determined by the initial parasitaemias (Figure 2B).

Trypanosomes were cultured as described in [38], in HMI9 with 10% fetal calf serum, starting with frozen mouse blood. For PAD1 staining, cells were methanol fixed, stained with anti-PAD1 antibody (gift from Keith Matthews, Edinburgh University) and DNA was counterstained with 1 μg ml-1 4′,6-diamidino-2-phenylindole, as described [20] except that the slides were incubated at 4°C overnight with the primary antibody. Stained slides were blinded before evaluation by an independent observer (CC); at least 100 cells were counted for each sample.

### Sequence analysis

Ribosomal RNA was depleted from the total RNA by hybridisation with antisense oligonucleotide and digestion with RNase H as described in [39].

Sequencing of genomic DNA and RNA was done using standard Illumina kits, and some genomic DNA was also sequenced using an Oxford nanopore device. Reads were aligned to the genome, then those aligning to open reading frames, annotated 3’-untranslated regions, and functional non-coding RNAs of the TREU927 genome were counted using a custom pipeline, tryprnaseq [40] [24]. Briefly, reads with trimmed using Cutadapt [41], then aligned using Bowtie2 [42] allowing for 2 mismatches. The transcriptome data were analysed using DeSeqU1 [43] [24], a custom version of DESeq2 [44].

To obtain gene copy numbers, all genomic DNA reads were allowed to align 20 times. This number was chosen because the vast majority of genes is present at less than 20 copies in the assembled TREU927 genome. We then selected unique genes, plus, for genes present in more than one copy, only a single representative copy, using a list slightly adapted from [29]. We then calculated reads per million per kilobase for this set of open reading frames, rounding to 2 decimal places. We next had to work out how many RPKM would represent a single-copy gene.

Since most genes in the core regions of the chromosomes are single-copy, we first calculated the modal RPKM value, expecting this to be near (though not precisely) the value for single-copy genes. (RPKM numbers were calculated to 2 decimal places and the counts for individual values were subject to considerable random variation.) We then adjusted the value slightly to get a symmetrical distribution for each dataset (Fig 4; S2 Table, sheet 7). We divided all RPKM values by this to attain gene copy numbers.

## Data availability

The genome sequences are available with accession numbers E-MTAB-9318 and E-MTAB-9759 for original strains, and E-MTAB-10457 and E-MTAB-10466 for cultures. The transcriptome results are deposited as E-MTAB-9320.

## Ethical approval

The sample collection and work with experimental animals was approved by the Makerere University Animal use committee. The rodent experiment approval is SBLS/HDRC/19/012.

## Acknowledgements

We thank Keith Matthews for antibody to PAD1, and George Cross and Annette McLeod for useful comments concerning the origin of the TREU927 genome DNA. JM is indebted to Prof. Michael Knop (ZMBH) for the loan of a fluorescence microscope. This work was partially funded by Deutsche Forschungsgemeinschaft grant number Cl112/28-1 to CC and JM.

## SUPPLEMENTS

### Supplementary text 1: Preliminary characterization

#### 1. *T. brucei* isolates from cattle grown in mice

We inoculated and successfully grew 3 recently isolated *Trypanosoma brucei brucei* strains from cattle (Table 1) and made fresh stabilates.

**Table 1.**

| Isolate ID | Simplified name | Source Village/Parish | Source District | Date of Isolation |
| --- | --- | --- | --- | --- |
| Tb065BAPC | MAK65 | Bunya | Apac | 01/02/2016 |
| Tb236BAPC |  | Bunya | Apac | 01/02/2016 |
| Tb098AAPC | MAK98 | Apuru | Apac | 30/07/2016 |

For simplicity Tb065BAPC and Tb098AAPC are designated MAK65 and MAK98 in the paper, emphasizing their original characterization at Makerere University. References in this text are at the end of the text.

#### 2. PCR characterization

##### Internal transcribed spacer (ITS)

We confirmed *Trypanozoon* status in all isolates by carrying out internal transcribed space (ITS) PCR, which yields a band size of approximately 480bp [1]. We confirmed that the parasites were *T. brucei brucei*, and therefore not infective for humans, by PCR for the *SRA* gene [2]. Human-infective *T. brucei rhodesiense* yield a product of 284bp. In both cases *T. b. rhodesiense* LW042 [3] was used as a positive control.

**Figure 1:**
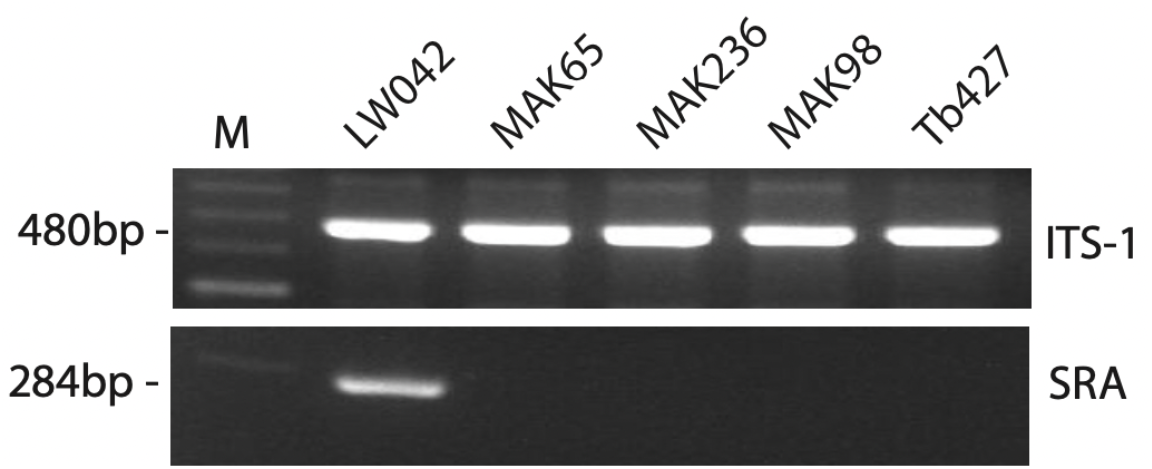
PCR confirmation that MAK65 and MAK98 are *T. brucei brucei*: positive PCR for ITS-1 and negative result for *SRA*.

##### Microsatellites

Genotyping of the *T. brucei* strains was carried out using microsatellites designated for *T. brucei* (Table 1) as described in [4].

The three recently isolated strains were again compared with *T. b. rhodesiense* human isolate LW042 (Figure 2). Tb236B and MAK65 were similar and could be distinguished from both MAK98 and LW042. This was probably because Tb236B and MAK65 were isolated from cattle in the same village/parish. Since we wanted to investigate different strains, Tb236B was not studied further.

**Figure 2:**
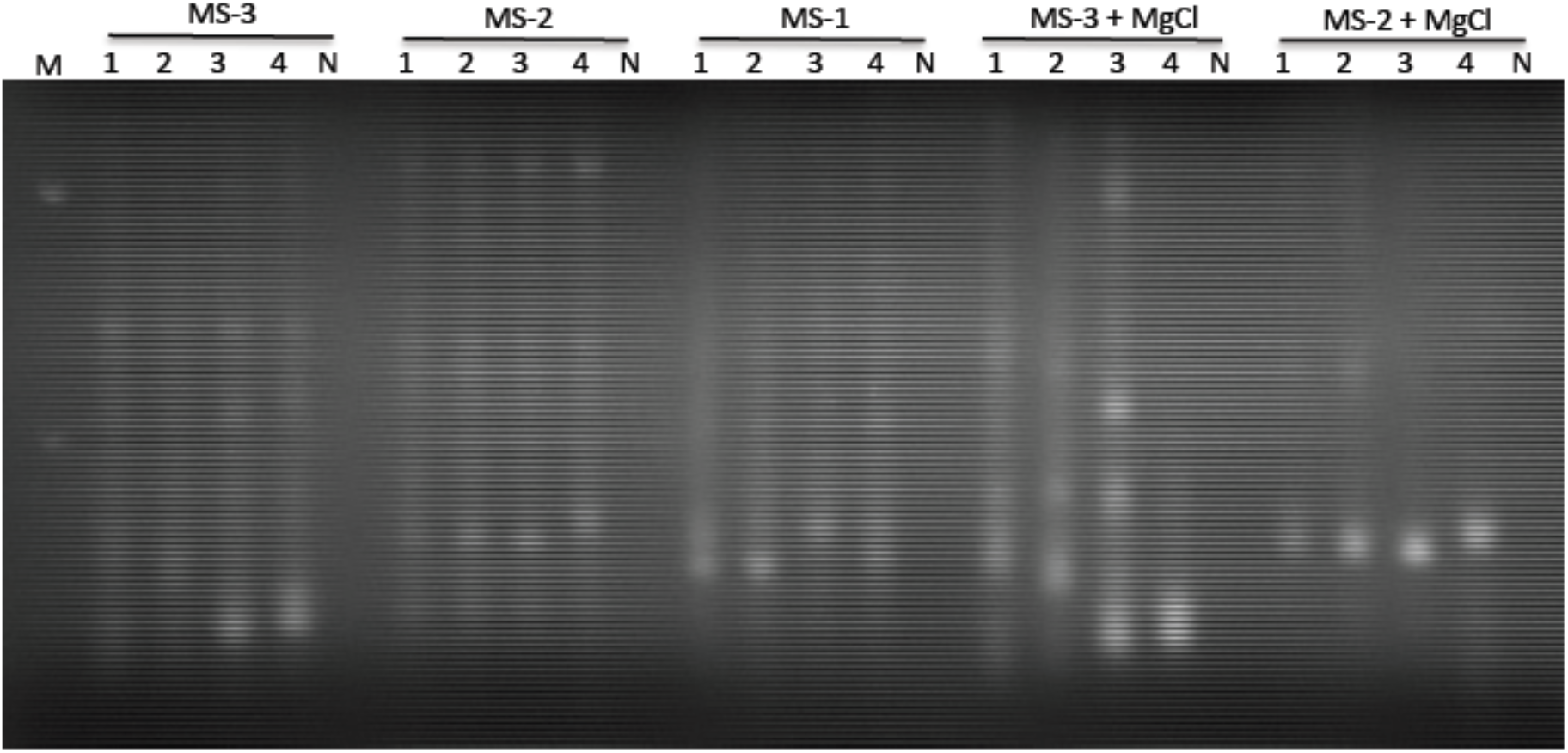
Microsatellite analysis.

Microsatellite analysis of 1-Tb236B, 2-MAK65, 3-LW042, 4-MAK98 using the primers shown in Table 2.

**Table 2.**
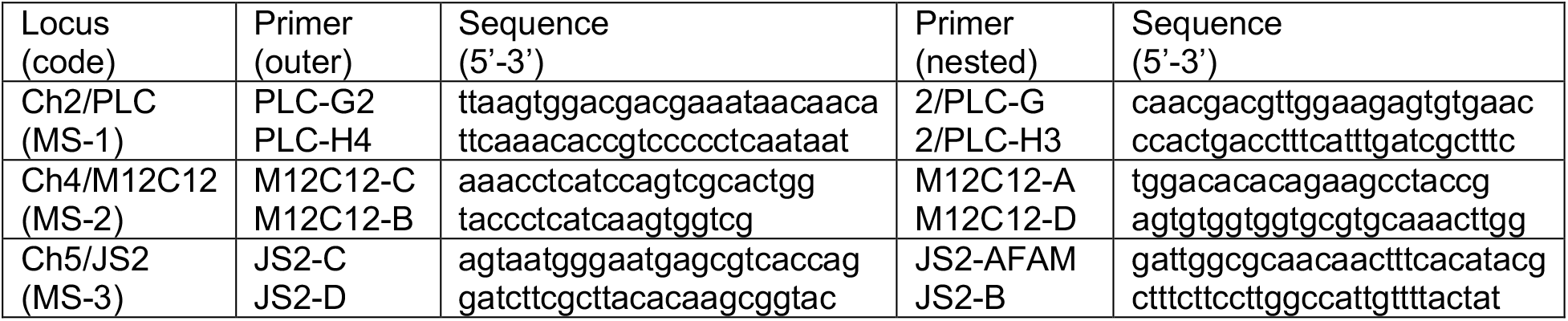

---

**S1 Figure.**
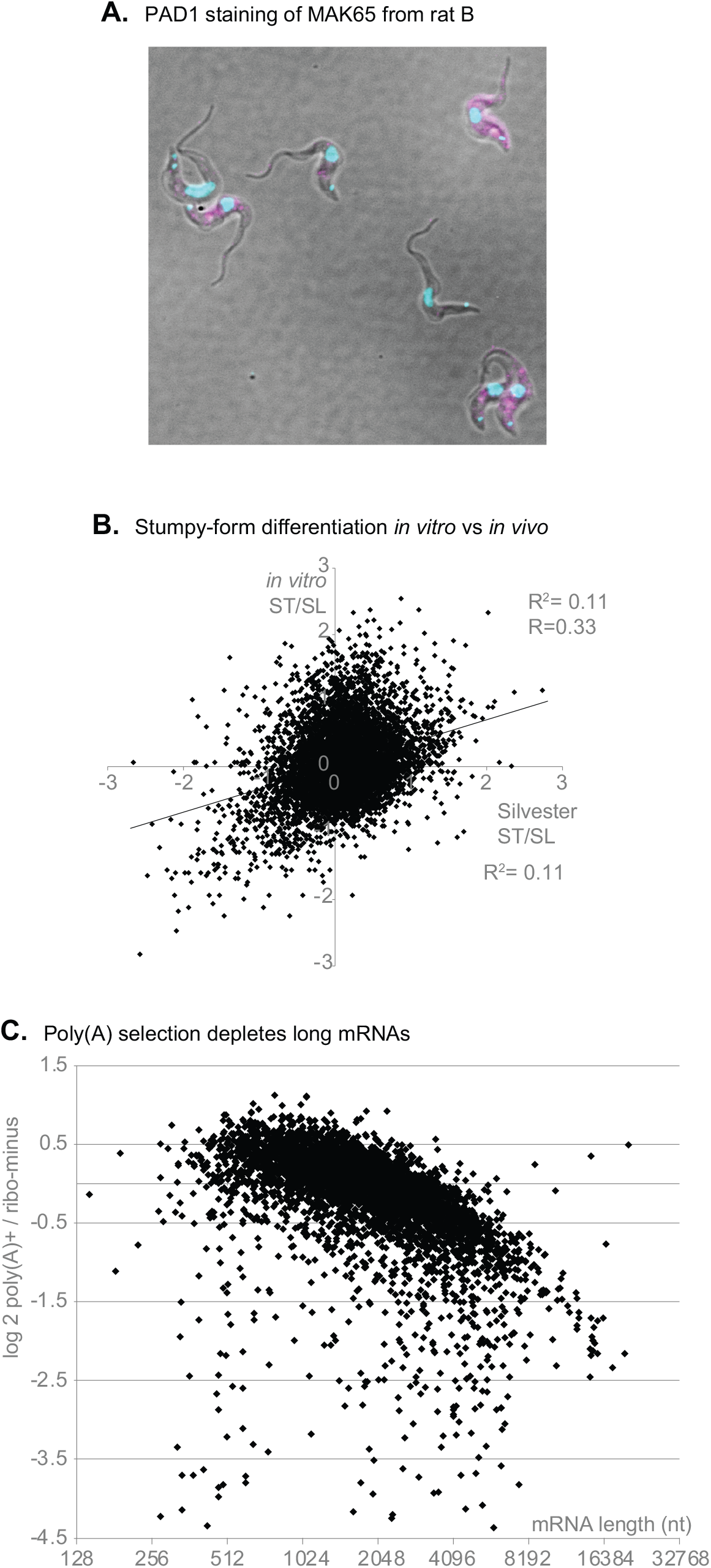
Transcriptome comparisons. **A**. Example of PAD1 staining for sample 65A. PAD1 is in magenta and DNA is cyan. **B**. Comparison of published results for *in vitro* [23] and *in vivo* [22] stumpy differentiation. In each case the log2 ratio of stumpy-form to long-slender form EATRO1125 is shown. **C**. Total RNA from MAK98 trypanosomes (sample from rat A) was either selected on oligo d(T) to give poly(A)+ RNA, or treated with RNase H and oligonucleotides complementary to the rRNA in order to give ribo-minus RNA. The log2 ratio of poly(A)+ to ribo-minus was is on the y-axis and the annotated mRNA length (log scale) on the x-axis. Results are in Supplementary table S1, sheets 2 and 5.

**S2 Figure.**
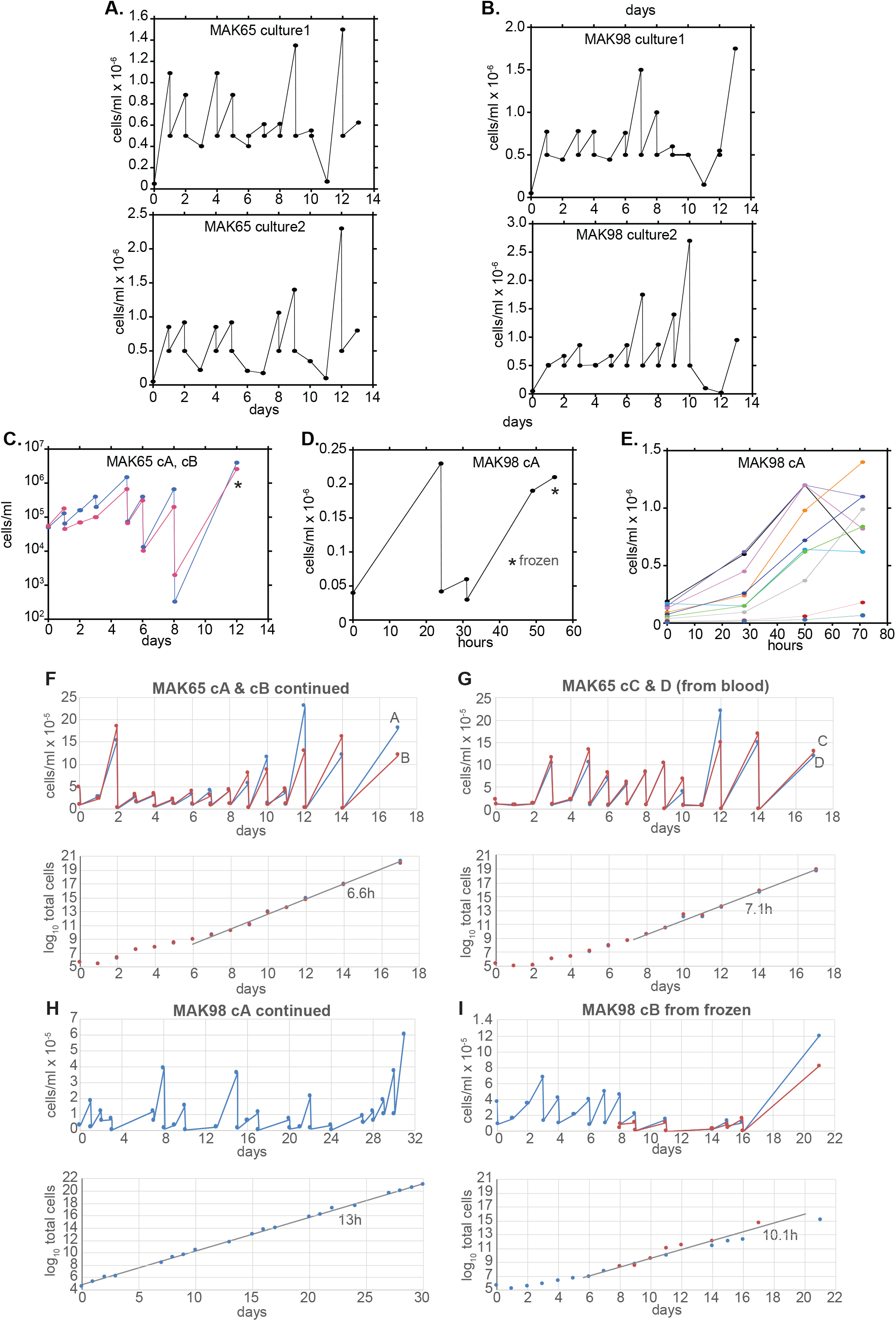
Cultures of MAK65 and MAK98 These data were used to make the cumulative curves in Figure 3; histories are in Figure 1. A. Initial cultures of MAK65, with parasite densities shown on a linear scale with dilutions B. Initial cultures of MAK98, with parasite densities shown on a linear scale with dilutions In each case the upper panel shows parasite densities on a linear scale and the lower panel shows cumulative parasite numbers on a log scale. The lines on the log scale graphs indicate the part used for division time calculations. In each case DNA was harvested at the end of the culture period, and stabilates were made. C. Cultures of MAK65 (cultures A and B) prior to creation of frozen stocks D. Culture of MAK98 (A) prior to creation of frozen stocks E. The MAK98 culture from (D) was subjected to serial dilutions in order to determine the maximum cell density. F. MAK65 cultures A and B (cA and cB) were continued from frozen stocks (panel C) to give a total culture time of 30 days. The upper panel is on a linear scale, and the lower panel is on a log scale showing the division time after 3 weeks of culture adaptation. G. MAK65 cultures C and D (cC and cD) were freshly initiated from blood stabilates. Details are as for (A) except that the culture time is shorter. H. Mak98 culture A (cA) was continued directly from panel D. I. Mak98 culture B (cB) was initiated from a frozen stabilate from Figure 2F. For genome analyses results from the two final cultures were >99% identical so the counts were pooled.

**S3 Figure.**
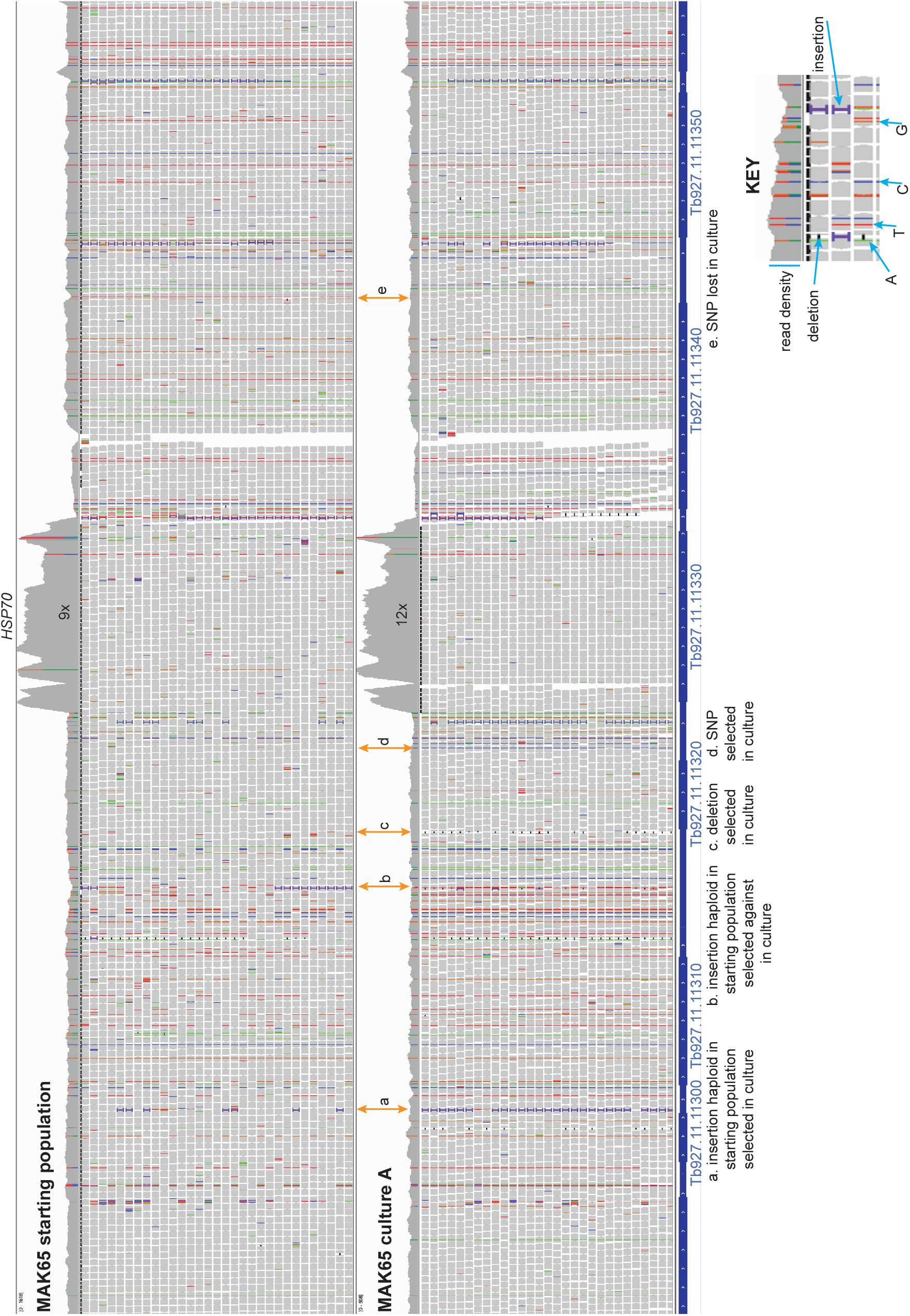
A detailed comparison over the *HSP70* locus for one MAK65 culture All reads for MAK65 starting population, and culture A, were allowed to align once to the TREU927 genome. The resulting mapped reads were visualized using the Integrated genome viewer (Broad Institute). The region surrounding the gene encoding the major cytosolic HSP70 is shown. The relative copy number for *HSP70* can be seen by comparing its read density with that over the surrounding single-copy regions. A key is below the alignment and few differences between the genomes are highlighted; these show regions where there might have been selection for particular variants. Analysis of more independent cultures would be needed to identify the most significant selective events.

**S1 Table**

Transcriptomes of MAK65 and MAK98 trypanosomes grown in rats. For details see the top sheet of the table.

**S2 Table**

Gene copy numbers for MAK65 and MAK98 trypanosomes grown in rats and in culture. For details see the top sheet of the table.

